# A *Caulobacter* hub taxon in the soybean root endosphere has a distinctive biofilm and engages in riboflavin-mediated mutualism with *Bradyrhizobium*

**DOI:** 10.64898/2026.09.14.750808

**Authors:** Madeline A. Quinlan, Sean Crosson, Sarah L. Lebeis, Gregory Bonito, Reid Longley, Aretha Fiebig

## Abstract

While it is well established that communities of microbes colonize the surfaces of plants as epiphytes and the interior plant tissues as endophytes, their ecological roles and colonization mechanisms remain under characterized. Members of the genus *Caulobacter* have emerged as hub taxa in epiphytic and endophytic microbiomes of diverse plant hosts, but the factors that support their colonization and community interactions have not been defined. Here, we characterize *Caulobacter* sp. RL271, a non-canonical member of the genus recently identified as a hub taxon in the soybean root endosphere. We demonstrate that the surface attachment strategies and biofilm architecture of *Caulobacter* sp. RL271 are determined by a matrix of capsular polysaccharide and cellulose rather than a polar holdfast adhesin, a classical defining feature of the genus. Both polysaccharide components influence the kinetics of plant root colonization in the model plant *Arabidopsis thaliana*. We further demonstrate that the ecology of *Caulobacter* sp. RL271 is shaped by its nutritional dependence on exogenous riboflavin and by interactions with *Bradyrhizobium diazoefficiens*, the nitrogen-fixing symbiont of soybean roots. This work advances understanding of *Caulobacter* biology and establishes RL271 as a tractable model for dissecting the functional role of hub taxa in root endophyte communities.

## Introduction

Interactions between rhizosphere microorganisms and their plant hosts are a major determinant of plant performance (Chepsergon and Moleleki, 2023). The diversity of microbes that inhabit rhizosphere communities is vast, but the alphaproteobacterial genus *Caulobacter* has emerged as a surprisingly consistent member of soil and plant-associated environments (Wilhelm, 2018). This ecological prevalence was not anticipated from the early history of the genus. Pioneering surveys by Poindexter isolated *Caulobacter* primarily from freshwater habitats (Poindexter, 1964, 1981), establishing its reputation as a freshwater oligotroph capable of persisting under extreme nutrient limitation. Culture-independent metagenomic surveys have since overturned this view, revealing that *Caulobacter* species are broadly distributed across both aquatic and terrestrial environments and are especially abundant where decaying plant material is available (Wilhelm, 2018).

Within plant-associated microbiomes, *Caulobacter* is more than an incidental colonist. *Caulobacter* was identified as a hub taxon in *Arabidopsis thaliana* (*Arabidopsis*) phyllosphere microbiomes (Agler et al., 2016), and has been repeatedly identified in and/or isolated from the endosphere and rhizosphere of roots of maize, soybean, and other hosts (Ulrich et al., 2008; Brown et al., 2012; Moya et al., 2017; Gao et al., 2018; Berrios and Ely, 2019; Luo et al., 2019; Yang et al., 2019; Gao et al., 2021; Ai et al., 2022; Bender et al., 2022; Hossain et al., 2023; Mejia et al., 2025; Qiao et al., 2026). Soybean is of particular interest because it is a major global crop and its productivity depends heavily on symbiotic nitrogen fixation by *Bradyrhizobium diazoefficiens*, which colonizes root nodules and provides the plant with fixed nitrogen (Peoples et al., 2021). *Caulobacter* sp. RL271 was recently identified as a hub taxon in the soybean root endosphere in a no-till agricultural field, placing it at the ecological center of a community that resides within a *Bradyrhizobium*-dependent host (Longley, 2022). Inoculation of a mixture of sand and field soil with a consortium of five hub taxa, including *Caulobacter* sp. RL271, enhanced soybean biomass under water limitation, increased root nodule counts, elevated expression of nodulation-associated plant genes, and boosted *Bradyrhizobium* levels (Longley, 2022). Strikingly, supplementation of the sand - field soil mix with *Caulobacter* sp. RL271 alone was sufficient to recapitulate the plant growth enhancement conferred by the five-taxa consortium. These observations raise the possibility that *Caulobacter* sp. RL271 influences soybean performance at least in part through direct effects on *Bradyrhizobium*, a hypothesis supported by a companion study demonstrating that *Caulobacter* sp. RL271 promotes *B. diazoefficiens* growth in vitro (Longley, 2022).

Understanding how *Caulobacter* sp. RL271 functions within its community requires understanding of its biology, but most of what is known about *Caulobacter* species is derived from studies of the freshwater isolate *C. crescentus* (Barrows and Goley, 2023). During its life cycle, *C. crescentus* differentiates from a motile swarmer cell to a stalked cell and produces polar structures that include a flagellum, pili, a stalk, and the holdfast adhesin. The holdfast is an exceptionally strong carbohydrate-based adhesive (Tsang et al., 2006; Berne et al., 2013), and genome-wide studies have identified a large regulatory and biosynthetic network that governs its assembly and function (Hershey et al., 2019a). However, comparative genomic analyses indicate that these surface-associated features, including the holdfast, are evolutionarily labile (Hallgren et al., 2025). This lability is part of a broader genomic and ecological divergence within the genus. Phylogenomic and metagenomic analyses indicate that terrestrial *Caulobacter* lineages are generally distinct from aquatic lineages and often have larger genomes, consistent with adaptation to different environmental demands (Wilhelm, 2018; Hallgren et al., 2025). Together, these results indicate that *Caulobacter* isolates like RL271 are not simply plant-adapted versions of *C. crescentus*, but members of distinct lineages with their own genomic repertoires, nutritional strategies, and surface-interaction traits.

One feature that may distinguish plant-associated *Caulobacter* is metabolic interdependence with neighboring microbes, particularly with respect to cofactor provisioning. In rhizosphere communities, flavins function not only as essential microbial cofactors but are secreted into the extracellular milieu. Rhizobia commonly release riboflavin (vitamin B2) or its derivatives into the surrounding environment (Phillips et al., 1999; Dakora et al., 2015), which impacts root colonization (Yurgel et al., 2014). Secreted flavins can also shape the broader rhizosphere microbiome assembly by supporting the growth of auxotrophs that cannot synthesize these cofactors themselves (Garrido-Sanz and Keel, 2025). Because *Caulobacter* sp. RL271 co-occurs with *Bradyrhizobium* in soybean root communities, it may be positioned to exploit this extracellular flavin pool.

A second axis of divergence in the genus is composition of the extracellular matrix. Surface colonization by *Caulobacter* is usually framed through the lens of holdfast-mediated adhesion (Merker and Smit, 1988; Smith et al., 2003; Hershey et al., 2019b). However, not all *Caulobacter* species produce a holdfast, and other extracellular polymers can contribute to surface attachment and biofilm architecture. For example, *C. crescentus* also produces a capsule-like exopolysaccharide that influences interactions with bacteriophage (Marks et al., 2010; Ardissone et al., 2014; Herr et al., 2018; McLaughlin et al., 2026), and impacts colony morphology (Marks et al., 2010) and biofilm structure (Fiebig, 2019). Bacterial cellulose is another major biofilm matrix polymer whose synthesis and export systems vary widely across bacteria, and that is commonly produced by plant-associated bacteria (Romling and Galperin, 2015; Abidi et al., 2022; Krasteva, 2024). It therefore plausible that plant-associated *Caulobacter* species will rely on a distinct suite of extracellular polymers, including cellulose, to support plant surface attachment and interactions with neighboring microbes.

Here, we characterize *Caulobacter* sp. RL271, a non-canonical member of the genus isolated from the soybean root endosphere. Using an integrated phylogenomic, genetic, and physiological approach, we examine its evolutionary placement within the family, define the extracellular matrix determinants that support biofilm formation and plant colonization in the absence of a canonical holdfast, and uncover ecological dependencies that are relevant to its root endosphere niche. This study establishes *Caulobacter sp.* RL271 as a tractable model for investigating the role of a hub taxon in root endophyte communities and supports a role for this genus as an ecological facilitator in plant associated communities.

## Results

### *Caulobacter* sp. RL271 has genome characteristics consistent with its terrestrial isolation

We sequenced the genome of *Caulobacter* sp. RL271 (hereafter RL271) using a combination of short and long read approaches. The assembly yielded a single circular chromosome of 5,687,505 bp with no detected plasmids. At 5.7 Mb, this is among the largest *Caulobacter* genomes on record, consistent with its terrestrial rather than aquatic origin (Wilhelm, 2018; Hallgren et al., 2025). To place RL271 in phylogenetic context, we constructed a maximum likelihood tree from single-copy core genes across 34 *Caulobacter* genomes spanning both aquatic and terrestrial isolation sites (Figure 1, Table S2) and compared average nucleotide identity (ANI) among these isolates (Figure S1). By both metrics, RL271 is most closely related to *Caulobacter* strain CBR1, a root isolate from an unidentified plant host (Berrios and Ely, 2019). These two strains share 91.8% ANI and form a distinct clade that is a sister group to *C. segnis* and *C. vibrioides* (more commonly known as *C. crescentus* and referred to as such hereafter). RL271 shares approximately 86% ANI with both *C. segnis* and *C. crescentus*, which is well below the 95% threshold conventionally used to delimit species boundaries, and therefore cannot be assigned to either described species.

**Figure 1.**
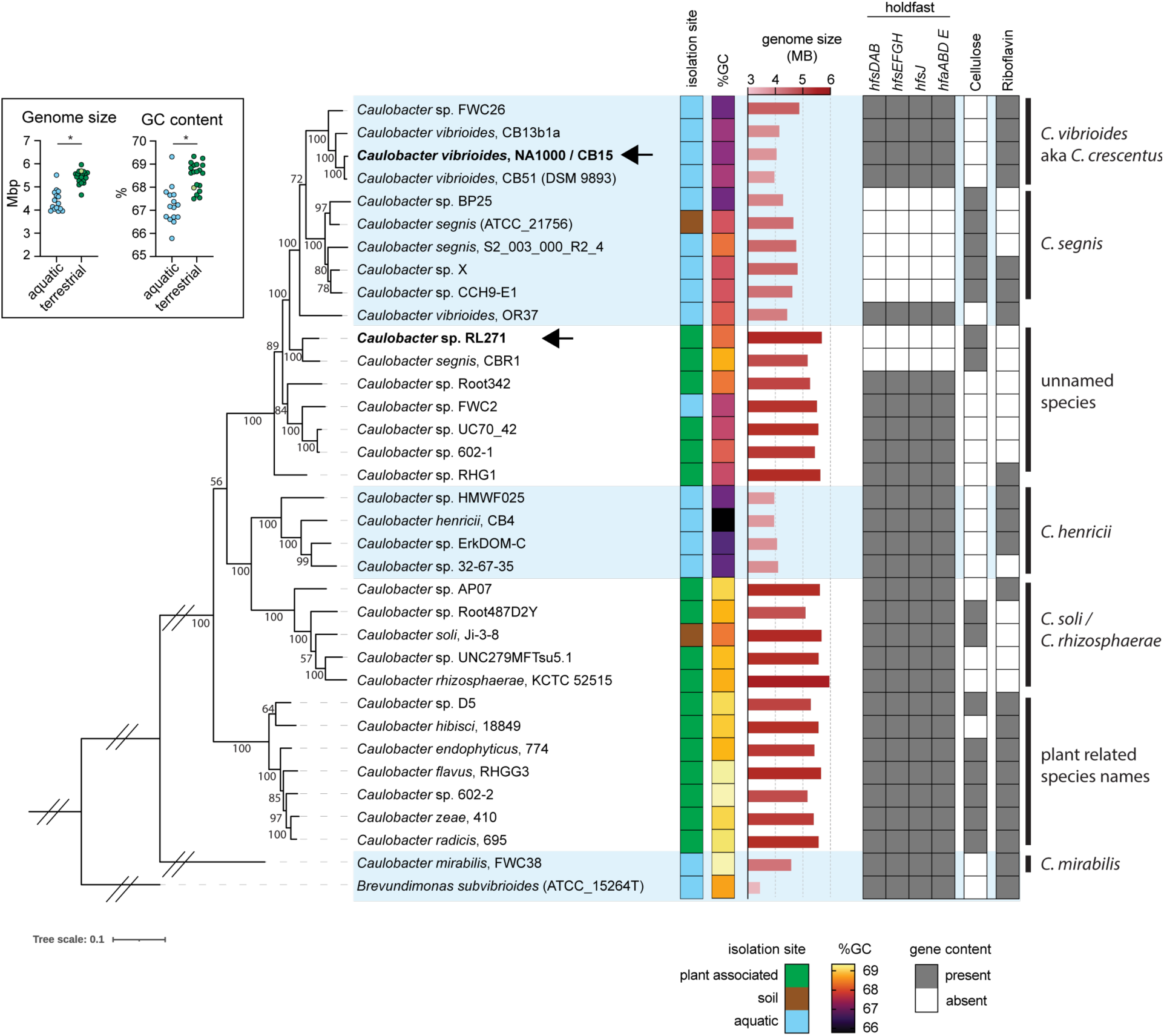
Phylogenetic and genomic features of *Caulobacter* sp. RL271 relative to other members of the *Caulobacter* genus. Maximum likelihood phylogenetic tree of 34 *Caulobacter* genomes rooted with *Brevundimonas subvibrioides* as an outgroup, constructed from 117 single-copy core genes with bootstrap support from 1,000 replicate trees. *Caulobacter* sp. RL271 and the model laboratory strains *C. vibrioides* (aka *C. crescentus*) NA1000/CB15 are indicated by arrows and bold type. Major species-level clades are labeled at right. Selected metadata are displayed as annotated columns alongside the tree. Environmental source is color-coded as follows: green, plant-associated (roots, leaves, endosphere, and rhizosphere); brown, soil (terrestrial isolates not described as plant-associated); blue, aquatic (freshwater sources including lakes, ponds, groundwater, wastewater, and built-environment plumbing). Percentage GC content is shown as a color gradient. Genome size is shown as a horizontal bar chart. Gene content columns indicate the presence (dark gray) or absence (white) of operons required for holdfast synthesis (*hfsDAB*, *hfsEFGH*, *hfsJ*) or anchoring (*hfaABDE*), cellulose synthesis (*bcsACDZ*), and riboflavin biosynthesis (*ribD-E-BA-H*). Genome size and GC content as a function of isolation site are plotted in inset in upper left. Groups were compared using a two-tailed, non-parametric Mann Whitney test (*, p<0.0001). GenBank accessions and full metadata for each isolate genome are provided in Table S2.

*Caulobacter* isolates from freshwater and terrestrial plant-associated habitats generally fall into distinct clades (Figure 1). Terrestrial plant-associated *Caulobacter* have larger genomes than aquatic species (Figure 1, inset; p<0.0001), consistent with broader comparative work showing that plant-associated and terrestrial bacteria often have larger genomes than related, non-terrestrial lineages (Levy et al., 2017). In our dataset, the terrestrial clades also show higher GC content (Figure 1, inset; p<0.0001). As aquatic and soil environments exist on a continuum and exchange of microorganisms between them occurs, a few exceptions to this pattern are expected. The *C. segnis* type strain (ATCC 21756), for instance, was isolated from soil yet clusters most closely with freshwater isolates and shares their genome characteristics (Urakami et al., 1990). Conversely, strain FWC2 was isolated from Lake Washington but has genome characteristics more typical of its plant-associated phylogenetic neighbors (Abraham et al., 1999). One interpretation of such cases is that aquatic *Caulobacter* populations are periodically replenished by water-borne dispersal from terrestrial habitats (Wilhelm, 2018). Nevertheless, the overall distributions of genome size and GC content across isolates suggest that aquatic and terrestrial *Caulobacter* populations are largely distinct, and that cross-habitat dispersal does not obscure the broader ecological signal. Moreover, the genome characteristics of RL271 are consistent with its isolation from soybean roots.

### Bradyrhizobium *rescues the riboflavin auxotrophy of* Caulobacter *sp. RL271*

To explore potential cross metabolite exchange within the endophytic soybean root microbiome, we performed metabolic modeling of the RL271 genome in KBase (Arkin et al., 2018), including gapfilling and flux balance analysis across media conditions, which predicted auxotrophies for riboflavin, spermidine, and asparagine. The predicted riboflavin auxotrophy has a clear genomic basis. RL271 lacks the four-gene operon (*ribD-ribE-ribBA-ribH*) required for riboflavin biosynthesis, as does the *C. segnis* type strain (ATCC 21756), a known riboflavin auxotroph (Urakami et al., 1990; Patel et al., 2015). Closely related isolates such as *Caulobacter* sp. X retain an intact *rib* operon, indicating that auxotrophy is not ancestral to this clade but has arisen independently (Figure 1, Table S2). More broadly, the riboflavin biosynthesis operon appears to have been lost and/or gained multiple times across the *Caulobacter* genus, with presumed auxotrophs distributed across both aquatic and terrestrial lineages (Figure 1, Table S2, (Hallgren et al., 2025)).

The consequences of riboflavin auxotrophy became apparent early in our work. RL271 was originally isolated on peptone-yeast extract (PYE) agar, but we observed inconsistent growth in PYE broth and traced this variability to age of the growth medium. As riboflavin is light-sensitive, we hypothesized that riboflavin degradation in broth stored in glass bottles under ambient light was responsible for the growth inconsistencies. To test this, we grew RL271 in freshly prepared PYE broth and in broth stored under ambient light for over one month, supplementing each with increasing riboflavin concentrations. Riboflavin supplementation had little effect on growth in fresh broth, indicating that this cofactor is not limiting at the time of preparation. In aged broth, however, growth was severely impaired and was restored by riboflavin supplementation in a dose-dependent manner, with maximal growth achieved at concentrations of 1 µM or above (Figure 2A). These results confirm that riboflavin degradation accounts for the variability in PYE broth performance. We therefore adopted the practice of using freshly prepared media or supplementing with 1 to 5 µM riboflavin at the time of inoculation for all subsequent experiments.

**Figure 2:**
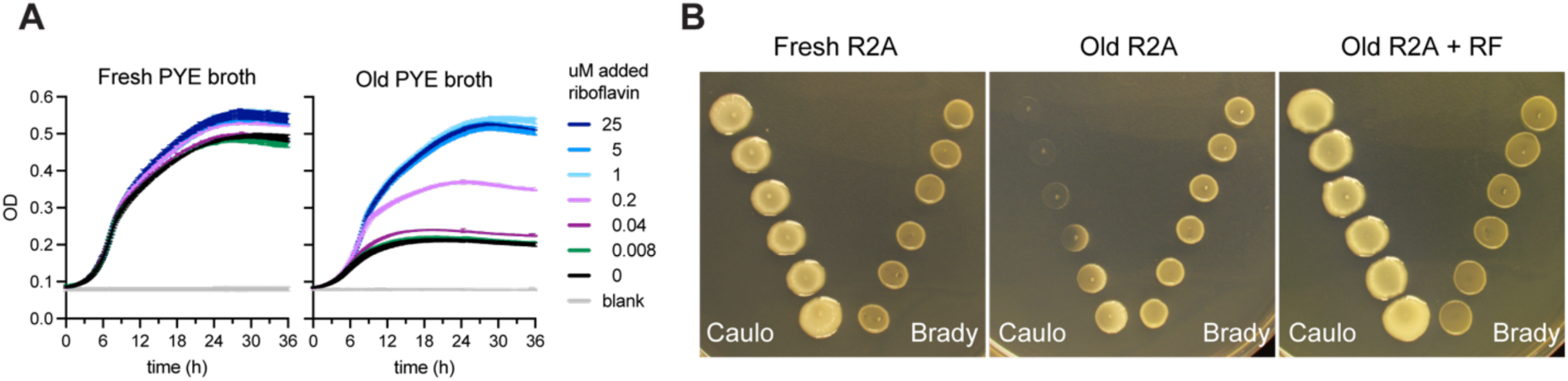
RL271 is a riboflavin auxotroph and *Bradyrhizobium* can compensate for this auxotrophy. **A)** Growth curves of wild-type RL271 in freshly made PYE broth (left) or PYE broth that had been stored at room temperature and room light for over a month (right). Each batch of media was supplemented with 0 to 25 µM riboflavin. Each line represents the mean and standard deviation of 3 technical replicates from a representative of three replicate experiments. **B)** *Caulobacter sp*. RL271 (left) and *Bradyrhizobium diazoefficiens* USDA110 (right) were spotted (5 µl per spot of cultures adjusted to 0.1 OD660) at increasing distances from each other on R2A agar that was freshly made (left), over 5 months old (center), or over 5 months old and supplemented with 5 µM riboflavin immediately before inoculation (right). Plates were imaged after 4 days at 30°C.

As mentioned, rhizobia are known to release riboflavin, which plays roles in plant root colonization and in modulating plant physiology (Dakora et al., 2015). Given that RL271 co-occurs with the nodulating soybean symbiont *Bradyrhizobium* in the soybean root endosphere, we hypothesized that secreted riboflavin from *Bradyrhizobium* could support RL271 growth under riboflavin-limiting conditions. To test this, we spotted RL271 at increasing distances from *B. diazoefficiens* USDA110 on fresh R2A agar, aged R2A agar, and aged R2A agar supplemented with 5 µM riboflavin. On freshly prepared medium, both species grew irrespective of their relative positions. On aged medium, RL271 growth was severely limited but was restored by proximity to *B. diazoefficiens*. Riboflavin supplementation prior to inoculation rescued RL271 growth regardless of its distance from *B. diazoefficiens*, demonstrating that the growth limitation on aged medium is attributable specifically to riboflavin deficiency (Figure 2B). These results support a model in which extracellular riboflavin released by *B. diazoefficiens* can cross-feed RL271 and mitigate its riboflavin auxotrophy.

### *Caulobacter* sp. RL271 retains dimorphic cell morphology but lacks holdfast

Like most *Caulobacter* strains, RL271 produces crescent-shaped cells and shows clear signatures of dimorphism (Figure 3A). In pre-divisional cells, the division plane is positioned asymmetrically, and motility is restricted to the smaller, newborn cells, consistent with the dimorphic life cycle described across the genus (Poindexter, 1981; Barrows and Goley, 2023; Hallgren et al., 2025; Hallgren and Jonas, 2025). However, RL271 produces little to no stalk, a trait shared with *C. segnis* and several species in the closely related genus *Brevundimonas* (Abraham et al., 1999; Patel et al., 2015; Curtis, 2017; Hallgren et al., 2025). The RL271 genome encodes genes for both the polar flagellum and type IV pili, which are elaborated on newborn swarmer cells in *C. crescentus*. In contrast, RL271 completely lacks the genes required for synthesis and cell-surface attachment of holdfast, a polar polysaccharide adhesin that mediates attachment of *C. crescentus* to diverse surfaces (Ong et al., 1990; Berne et al., 2013; Fiebig, 2019) and is otherwise a defining feature of the genus (Gupta and Mok, 2007; Wilhelm, 2018).

**Figure 3:**
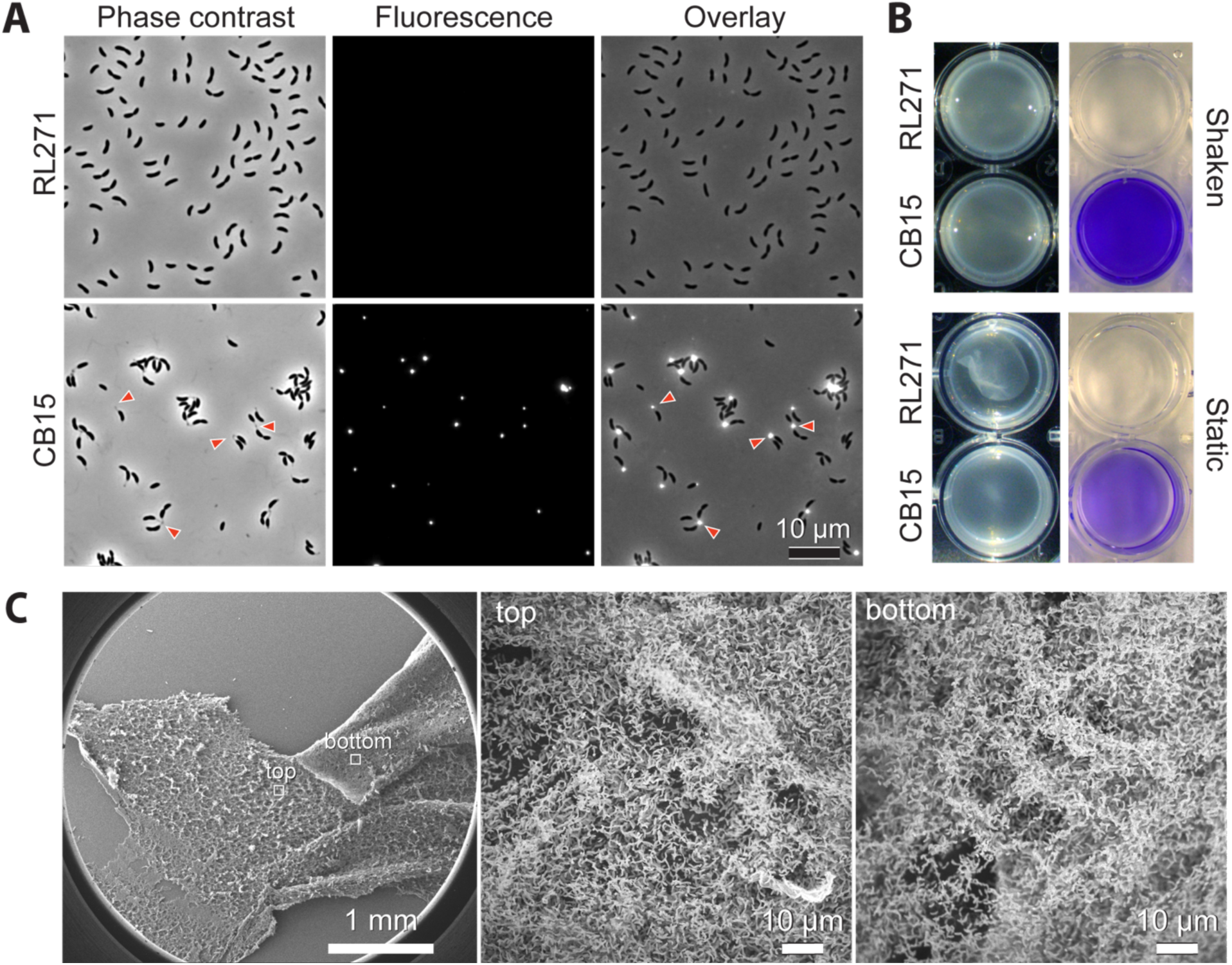
*Caulobacter sp.* RL271 biofilms do not attach to glass or plastic surfaces. **(A)** Light micrographs of *C. crescentus* CB15 and *Caulobacter sp.* RL271 stained with fluorescent WGA to label holdfast polysaccharide. **(B)** Overnight cultures of the same strains as in A grown in 24-well plates with shaking (top) or static without shaking (bottom). In each growth conditions, images of cultures before staining (left) are paired with images of the same wells after crystal violet staining of surface attached cells after the cultures were washed away (right). A floating biofilm that is washed away during staining can be observed at the bottom of the static RL271 culture. **(C)** Scanning electron micrographs of a RL271 biofilm loosely attached to a glass coverslip. *Left*: 25X magnification. *Middle*: top of film at 1000X, *Right*: underside of peeled film at 1000X magnification. Boxes on the low magnification image on the left indicate the approximate size of the higher magnification images to the right.

Consistent with this genomic absence, RL271 does not form rosettes, does not stain with holdfast-binding lectins (Figure 3A), and shows no appreciable adhesion to plastic surfaces by crystal violet staining (Figure 3B) or to glass surfaces by SEM (Figure 3C). Loss of holdfast biosynthetic capacity has occurred in a small number of other *Caulobacter* isolates, including strain CBR1 (Berrios, 2021), and *C. segnis* and its closest relatives (Patel et al., 2015; Hallgren et al., 2025), (Figure 1, Table S2).

### A genome-wide fitness screen reveals essential functions and unexpected growth dependencies in *Caulobacter* sp. RL271

To interrogate gene function and assess gene essentiality in the RL271 genome, we constructed a barcoded Himar transposon mutant library using a previously described system (Wetmore et al., 2015). The *Tn*-Himar insertion mutants were selected on PYE complex medium, and the library contained approximately 10^5^ clones. Bulk mapping identified insertions at 52,302 of 68,267 TA dinucleotides (76.6%). In total, 59,076 barcodes mapped confidently to insertion sites, corresponding to a mean of 10 and a median of 7 barcoded strains per protein-coding gene.

We used two complementary approaches to predict genes whose disruption was strongly underrepresented in the mutant pool, i.e. candidate “essential” genes (DeJesus et al., 2015; Ioerger, 2022). A total of 301 genes were classified as essential or growth defective by both approaches; 442 genes were classified as essential or growth defective by at least one approach (Table S3). As expected, the high-confidence essential set included genes involved in chromosome replication and segregation (e.g., *dnaA*, *dnaN*, *dnaQ*, *gyrA*, *gyrB*, *parC*, *parE*), transcription (*rpoB*, *rpoC*, *rpoD*), cell envelope biogenesis (*bamA*, *lptB*, *lptE*, *murB*, *murC*, *murF*, *murG*, *mraY*, *murJ*), and cell division/cell-cycle control (*ctrA*, *ftsZ*, *ftsW*, *rodA*, *mreC*), consistent with prior *Caulobacter* essentiality analyses (Christen et al., 2011; Hentchel et al., 2019). Several additional high-confidence hits pointed to a strong dependence on aerobic energy metabolism and nutrient scavenging, including respiratory genes (*nuo* and *sdh* operons), the high-affinity phosphate transporter *pstC* (Lubin et al., 2016), and the *tolB-tonB-exbD-tolQ* membrane transport system.

One unexpected pattern was a cluster of predicted extracellular polysaccharide genes, in which 8 of 10 genes in the MZV50_RS16475–RS16520 region were classified as essential or growth defective (Table S3, Figure 4A). This locus encodes several predicted glycosyltransferases, acyltransferases, and hypothetical proteins. In addition, MZV50_RS25000, MZV50_RS25005, MZV50_RS25030, and MZV50_RS25035, which are homologous to genes required for capsule export in *C. crescentus* NA1000 (Ardissone et al., 2014), were also classified as essential or growth defective. Together, these results indicate that envelope polysaccharide biosynthesis impacts RL271 fitness under the conditions tested. The dataset further revealed an unexpected requirement for aromatic amino acid/tryptophan biosynthesis. This was surprising because classical *C. crescentus* genetic studies identified recoverable *trp* auxotrophs, including a *trpB* mutant (Ross and Winkler, 1988), indicating that tryptophan synthase is not uniformly essential in *Caulobacter*. The identification of glutamine synthetase (*glnA*) as an essential gene indicates that RL271 fitness is also unusually sensitive to nitrogen assimilation and glutamate/glutamine homeostasis, processes that have been tied to growth and cell-cycle control in *C. crescentus* (Beaufay et al., 2015; North et al., 2025).

**Figure 4.**
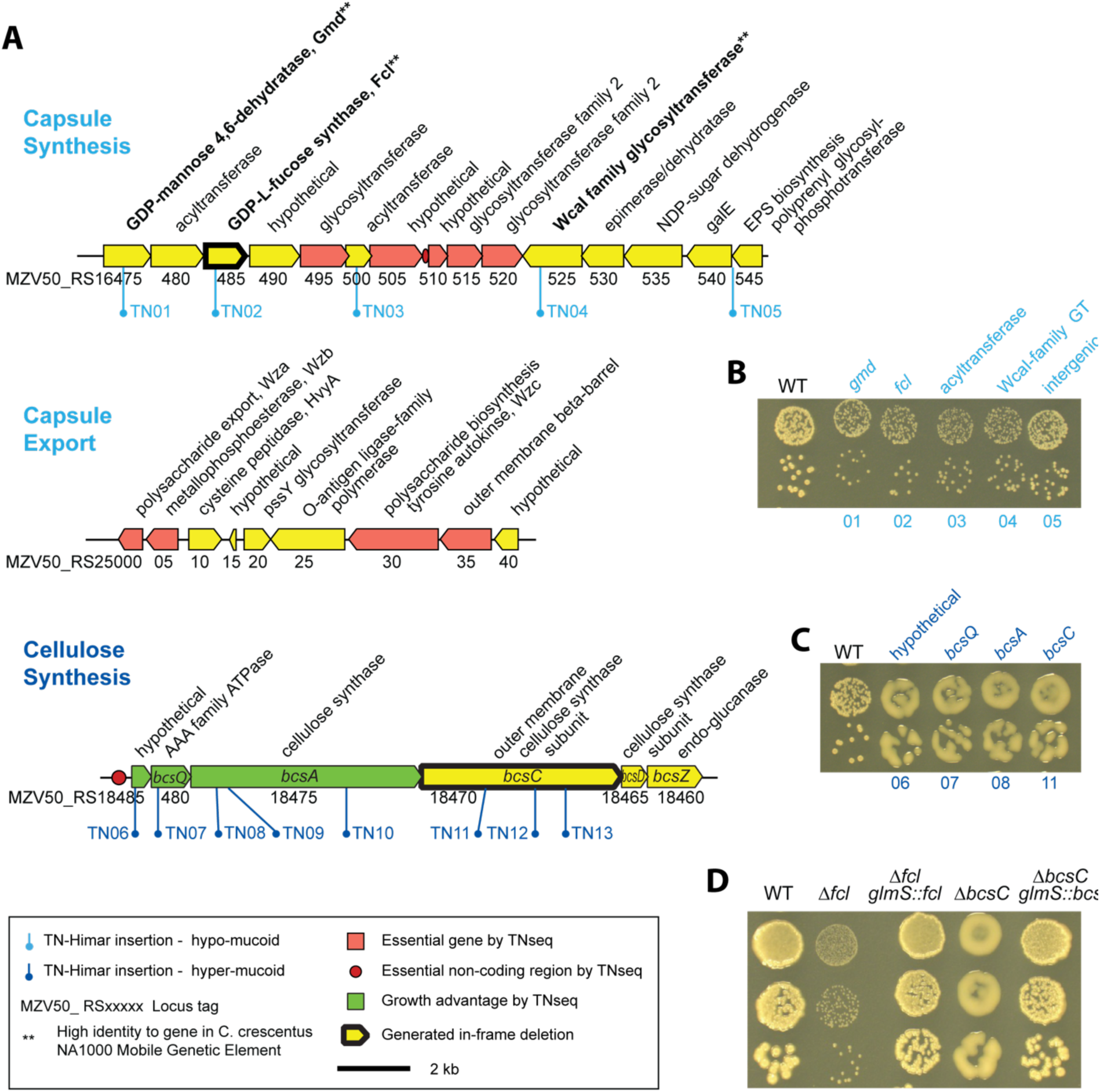
Genomic loci for capsule and cellulose biosynthesis and their contributions to colony morphology. **(A)** Schematic of three *Caulobacter* sp. RL271 genomic loci involved in extracellular polysaccharide biosynthesis and export: the capsule synthesis locus (MZV50_RS16475–RS16545), the capsule export locus (MZV50_RS25000 – RS25040), and the cellulose synthesis locus (MZV50_RS18460–RS18485). Gene colors indicate fitness classification from *Tn*-Himar analysis, and lollipop symbols indicate the positions of transposon insertions recovered from hypo-mucoid (light blue; TN01-TN05) or hyper-mucoid (dark blue, TN06-TN13) colonies identified. Chromosomal positions of the insertion sites in each strain are in Table S1. Asterisks indicate genes with high amino acid identity to genes in the *C. crescentus* NA1000 mobile genetic element. **(B-D)** Colonies from serially diluted cultures grown on R2A for 2 days at 30°C; **(B)** hypo-mucoid *Tn*-Himar insertion strains, **(C)** representative hyper-mucoid Tn-himar insertion strains, or **(D)** in-frame deletion and corresponding complementation strains. For transposon mutants, gene disrupted is above and clone number is below.

Beyond essential genes, HMM-based analysis (Ioerger, 2022) identified genes whose disruption conferred an apparent growth advantage under our culture conditions; ten RL271 genes fell into this category. Three of these lie within a predicted cellulose biosynthesis operon (Table S3, Figure 4A). Although experimental evidence for cellulose synthesis in *Caulobacter* has not previously been reported, bacterial cellulose is widespread among plant-associated bacteria, where it commonly contributes to surface attachment and biofilm formation (Matthysse et al., 2005; Thompson et al., 2018; Heredia-Ponce et al., 2021). The RL271 locus most closely resembles a group 1b cellulose synthase system as defined by Römling and Galperin (Romling and Galperin, 2015). Like other type 1 systems, it encodes the cellulose synthase BcsA, the export factor BcsC, and BcsD, which promotes crystalline cellulose formation. Its classification as type 1b reflects the additional presence of *bcsQ*, a polarly-localized cytoplasmic AAA ATPase whose precise role remains unresolved but is thought to localize synthesis and/or secretion of cellulose (Le Quere and Ghigo, 2009; Romling and Galperin, 2015; Abidi et al., 2022; Krasteva, 2024). Inspection of other *Caulobacter* genomes revealed that this operon is present in strain CBR1 and other members of the *C. segnis* group. A related operon, lacking *bcsQ* and carrying a distinct hypothetical accessory gene, was found in some, but not all, soil- and root-associated species, including *C. soli*, *C. endophyticus*, and *C. zeae* (Figure 1, Figure S2). Collectively, analysis of this rich mutant collection revealed expected essential genes, unexpected requirements for amino acid biosynthesis, and unexpected fitness impacts for extracellular polysaccharide biosynthesis in RL271.

### *Caulobacter* sp. RL271 forms structured biofilms despite lacking holdfast

Biofilm formation confers many protective advantages and can support surface colonization, such as the growth of bacteria directly on plant roots (Flemming et al., 2016; Knights et al., 2021). Given that RL271 lacks the genetic capacity to produce holdfast, the defining surface adhesin of the *Caulobacter* genus, we sought to determine what mechanisms it uses for surface attachment and biofilm formation. First, we simply inspected RL271 cultivated under different conditions. RL271 formed aggregated cell clumps and larger flocs in shaken culture, while static growth conditions supported the formation of an organized film (Figure 3B), providing evidence for distinct adhesion mechanisms in RL271. Scanning electron microscopy of cells grown under static conditions revealed a cohesive, well-structured mat encased in a net-like fibrous material. Although the cells formed a structurally integrated biofilm, it showed little affinity for the underlying glass substrate and could be observed peeling away from the surface (Figure 3C), consistent with strong cell-cell but weak cell-surface interactions.

To identify the matrix components responsible for biofilm cohesion, we visually screened our transposon library for mutants with altered colony morphology, an approach that has successfully identified extracellular polysaccharide biosynthesis genes in other organisms (Kearns et al., 2005; Marks et al., 2010; Cabeen et al., 2016; Thompson et al., 2018; Onyeziri et al., 2022; Goetsch et al., 2024). We recovered mutants forming either smaller, hypo-mucoid or larger, hyper-mucoid colonies and mapped the transposon insertion sites in each class.

### Hypo-mucoid mutants reveal capsule polysaccharide biosynthesis genes

Insertions in small, hypo-mucoid (rough) colonies mapped to several gene classes, including three encoding Gmd (GDP-mannose 4,6-dehydratase; MZV50_RS16475), Fcl (GDP-L-fucose synthase; MZV50_RS16485), and a WcaI-family glycosyltransferase (MZV50_RS16525) (Figure 4A-B). Gmd and Fcl typically function together to convert GDP-D-mannose to GDP-L-fucose, a sugar nucleotide incorporated into surface polysaccharides such as colanic acid, capsule, and O-antigen (Andrianopoulos et al., 1998; Whitfield, 2006); in *E. coli* WcaI incorporates fucose into colanic acid (Scott et al., 2019). All three of these RL271 genes reside in a predicted polysaccharide biosynthesis cluster (Figure 4) and share high amino acid identity with genes in a 26 Kb mobile genetic element (MGE) of *C. crescentus* NA1000: CCNA_00472 (Gmd, 96%), CCNA_00471 (Fcl, 80%), and CCNA_03998 (glycosyltransferase, 54%). In NA1000, this MGE enables capsule polysaccharide synthesis, a function that requires all three genes (Marks et al., 2010; Ardissone et al., 2014; Herr et al., 2018). Although the flanking genes in the NA1000 MGE and this RL271 locus share little sequence similarity, they share predicted functional roles in polysaccharide biosynthesis. As in NA1000 (Christen et al., 2011), several genes at this locus are predicted to be essential by transposon mutagenesis (Figure 4, Table S3). By analogy with the NA1000 MGE, this apparent essentiality may reflect synthetic toxicity from accumulation of biosynthetic intermediates or competition for shared substrates rather than an absolute requirement for growth (Ardissone et al., 2014). A second locus encoding predicted capsule export genes in NA1000 (Ardissone et al., 2014) is conserved in RL271, and many of these genes were also classified as essential under our tested conditions (Figure 4, Table S3).

To confirm that disruption of these genes affects colony morphology, we constructed an in-frame deletion of *fcl* (Δ*fcl*). Indeed, colonies of this mutant are small and less mucoid, and this defect can be restored by integrating *fcl* at the *glmS* site (Figure 4D). To evaluate the role of *fcl* in capsule production, we used a fluorescent dextran exclusion assay to detect capsule around individual cells (Ardissone et al., 2014). Wild-type RL271 cells exclude dextran from a zone extending beyond the cell body, consistent with an elaborated capsule (Figure 5A top). This exclusion is variable across cells (Figure 5B), likely reflecting cell-cycle-dependent regulation analogous to the HvyA-mediated inhibition of capsule synthesis during G1 in *C. crescentus* NA1000 (Ardissone et al., 2014). Indeed, RL271 encodes a HvyA homolog (MZV50_RS25010) adjacent to the predicted capsule export genes (Figure 4). Deletion of *fcl* abolished dextran exclusion entirely (Figure 5A-middle, and 5B), and complementation by integrating *fcl* at the *glmS* site fully restored it (Figure 5B). We conclude that RL271 elaborates a capsule and that *fcl* is required for capsule polysaccharide synthesis.

**Figure 5:**
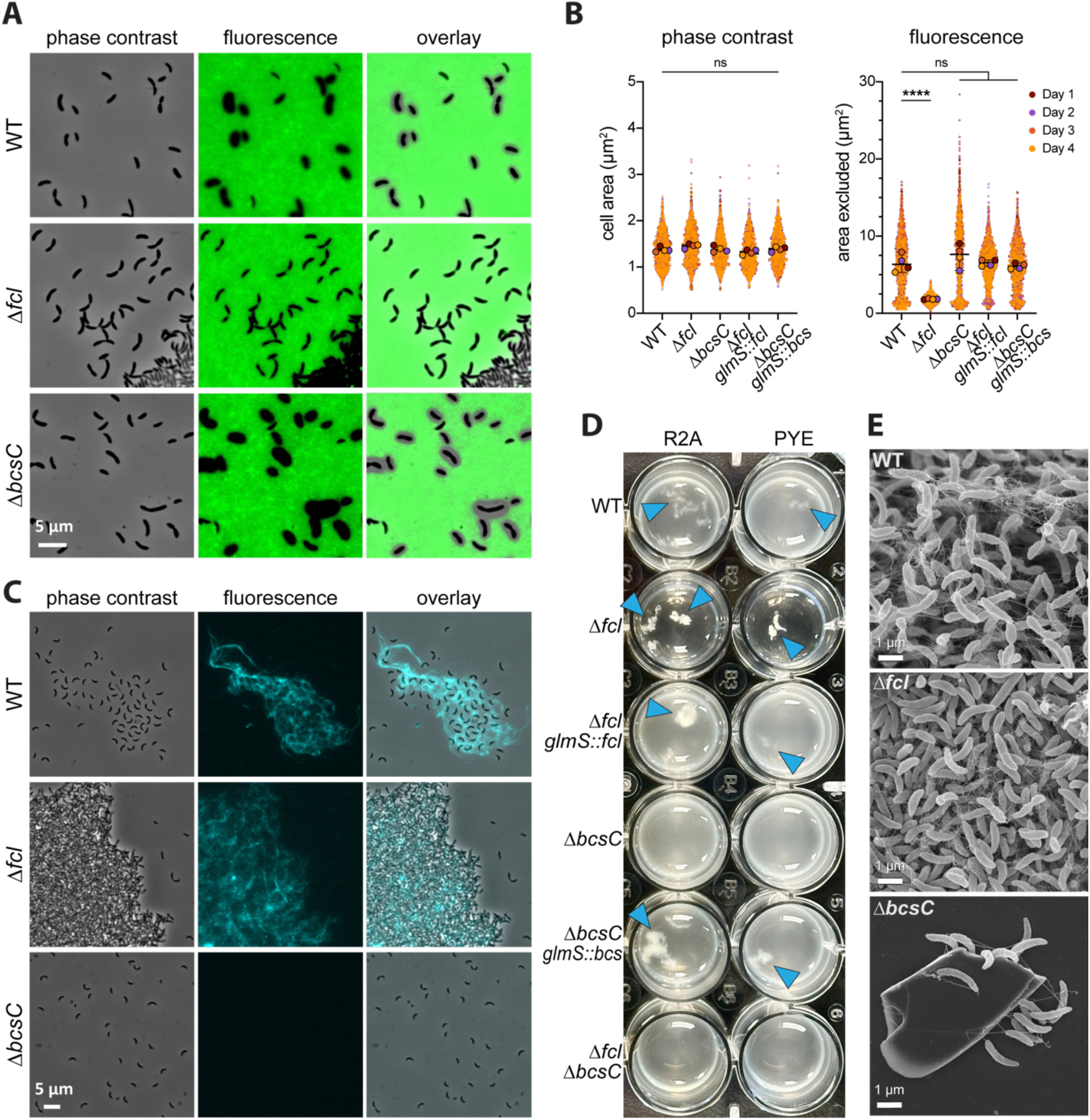
*Caulobacter* sp. RL271 elaborates an *fcl*-dependent capsule and cellulose, both of which contribute to biofilm architecture. **(A)** Negative stain reveals capsule surrounding RL271 strains. Representative phase contrast and fluorescence images of log phase cells grown in PYE broth incubated with FITC-dextran. The overlay image shows the dextran excluded areas around cell bodies. **(B)** Area of cell bodies (phase) and dextran excluded zones (fluorescence) for four independent cultures (at least 500 cells quantified per replicate) presented as super-plots (Lord et al., 2020). Small dots indicate individual cells (colored per replicate). The four large dots per strain represent the mean of each replicate. These summary values were compared to WT using a one-way ANOVA, followed by Dunnett’s post-test (ns, p >0.05; **** p < 0.0001). **(C)** Representative phase contrast and fluorescence micrographs of cells stained with 0.01% calcofluor white. Cells were grown on PYE agar plates and resuspended from plates for staining. The genotype corresponding to each set of images is indicated on the left. **(D)** Indicated strains cultured in R2A (left) or PYE (right) broth in 24-well plates with shaking for 24 hours. Blue arrowheads highlight flocks in cultures. **(E)** Scanning electron micrographs of biofilms of WT, Δ*fcl* and Δ*bcsC* mutants grown in static cultures taken at 10,000X magnification.

### Hyper-mucoid mutants reveal cellulose biosynthesis genes

Insertions in hyper-mucoid colonies mapped to the cellulose synthase locus, including a small hypothetical gene at the 5’ end of the operon (MZV50_RS18485), *bcsQ* (MZV50_RS18480), *bcsA* (MZV50_RS18475), and *bcsC* (MZV50_RS18470) (Figure 4A & 4C). As noted above, the first three genes in this operon were categorized as conferring a growth advantage when disrupted in our pooled *Tn*-Himar library, suggesting a penalty for cellulose synthesis under the conditions used to generate the library. Consistent with this, we recovered more mutant strains with disruptions in *bcs* genes than in capsule biosynthesis genes.

To validate the impact of cellulose locus mutation on colony morphology and matrix production, we generated an in-frame deletion of *bcsC* (Δ*bcsC*). Deletion of *bcsC* recapitulated the mucoid colony morphology phenotype of the *bcs* transposon mutants and the wild-type colony morphology was restored by expressing the *bcs* operon (MZV50_RS18485-60) from its native promoter inserted at the *glmS* site (Figure 4D). Dextran exclusion around Δ*bcsC* cells was not statistically different from wild type (Figure 5B). To assess cellulose production directly, we stained liquid cultures with calcofluor white, which binds β-glucan polymers including cellulose (Haigler et al., 1980). Fluorescent strands were observed surrounding clumps of wild-type cells, consistent with extracellular cellulose fibers (Figure 5C). These fibers were absent from Δ*bcsC* cells, confirming that *bcsC* is required for their production. In contrast, Δ*fcl* cells formed densely packed aggregates with robust calcofluor staining.

### Cellulose and capsule make distinct contributions to RL271 biofilm architecture

Finally, we used our deletion strains to dissect the contributions of each polysaccharide to biofilm formation. When shaken, the flocs observed in wild type cultures were absent in Δ*bcsC* cultures and restored by genetic complementation (Figure 5D). SEM confirmed that Δ*bcsC* cells failed to form cohesive mats. In rare cases, individual cells were observed attached to glass fragments, apparently via their flagella (Figure 5E). In contrast, Δ*fcl* cultures were nearly devoid of planktonic turbidity, as this mutant aggregated into large, stable flocs. Genetic complementation with *fcl* restored wild-type turbidity and floc size. SEM of Δ*fcl* biofilms revealed more densely packed cells than wild type, still encased in net-like fibrous material. To determine whether the hyperflocculant phenotype of Δ*fcl* mutants was cellulose-dependent, we constructed a Δ*fcl* Δ*bcsC* double mutant. These cultures were evenly dispersed, resembling the Δ*bcsC* single mutant, indicating that the dense flocs formed by Δ*fcl* cells require cellulose (Figure 5D). We conclude that cellulose is the primary structural determinant of RL271 biofilm cohesion, and that capsule modulates biofilm architecture by limiting close cell-cell packing and maintaining a more open, dispersed biofilm structure.

### Capsule promotes directed growth of Caulobacter sp. RL271 toward B. diazoefficiens

We have demonstrated that RL271 enhances the growth of *B. diazoefficiens* on soil extract plates (Longley, 2022). To examine whether either cellulose or capsule extracellular polysaccharides contribute to the growth enhancement of *B. diazoefficiens*, we spotted wild-type RL271, Δ*fcl* or Δ*bcsC* mutants adjacent to spots of *B. diazoefficiens* on soil extract plates. As expected, proximity to RL271 enhanced growth of *B. diazoefficiens*. We observed similar enhancement by both mutants and concluded that elaboration of the capsule and cellulose polysaccharides does not contribute to the impact of RL271 on *B. diazoefficiens* (Figure 6). In the spot assay described above, RL271 colonies do not simply grow in place but over time extended directionally toward *B. diazoefficiens*. As the RL271 colony expands beyond the region of initial inoculation, it becomes noticeably more translucent and mucoid, consistent with active capsule production at the leading edge. This directed growth was absent in the Δ*fcl* mutant, which lacks capsule (Figure 6). Together, these observations support a model in which the RL271 capsule acts as a hygroscopic matrix that facilitates directed movement toward neighboring microorganisms (in this case toward *B. diazoefficiens*) which we have shown can supply the riboflavin that RL271 cannot synthesize itself (Figure 2).

**Figure 6:**
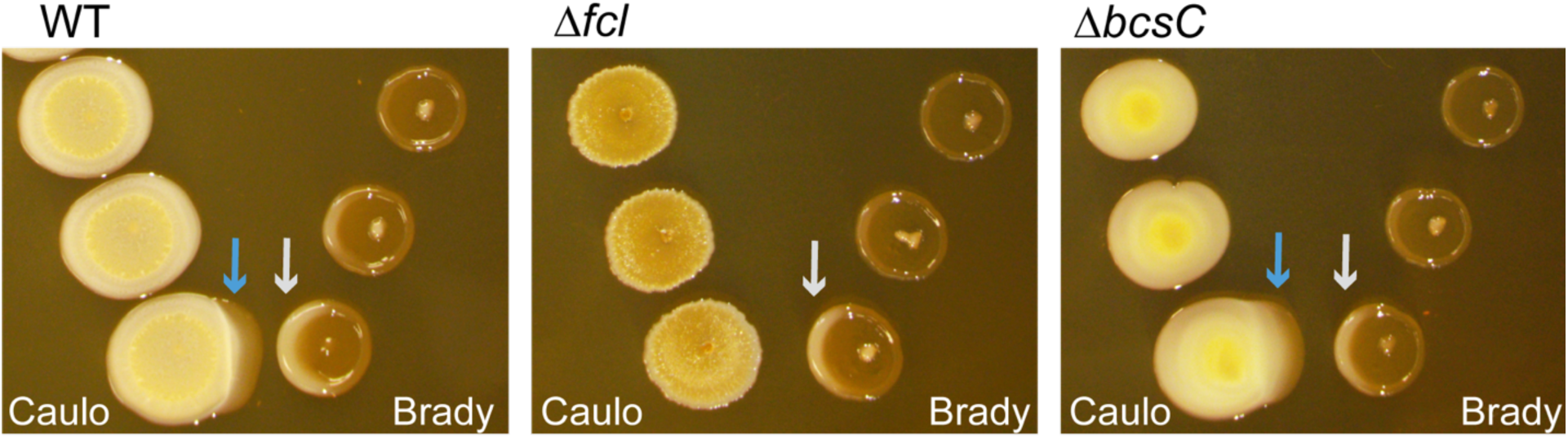
*Caulobacter* sp. RL271 migrates toward *Bradyrhizobium* in a capsule-dependent fashion. RL271 strains (left) spotted at increasing distances from *B. diazoefficiens* USDA 110 (right) on soil extract plates with xylose as the carbon source after 5 days of growth. The RL271 genotype is indicated above each image. The blue arrows highlight the directed migration of the RL271 spot colony toward *B. diazoefficiens*. The light grey arrows highlight the growth enhancement of *B. diazoefficiens* near RL271.

### Capsule and cellulose polysaccharides support colonization of plant roots by Caulobacter sp.RL271

To test whether either extracellular polysaccharide influences colonization of plant roots, we inoculated *Arabidopsis* seedlings grown under defined sterile conditions with wild-type RL271, Δ*fcl*, Δ*bcsC*, or Δ*fcl* Δ*bcsC* strains. At time points following inoculation, roots were harvested, homogenized, and titered to quantify bacterial load. RL271 did not grow detectably in this system in the absence of roots, confirming that root exudates provide essential nutrients required to support growth under these conditions.

Wild-type RL271 colonized roots efficiently across all time points, with populations increasing rapidly in the first 24 to 48 hours before plateauing (Figure 7A-B). All mutants reached densities comparable to wild-type by 2 to 4 days post-inoculation, indicating that neither polysaccharide is required for long-term root association. However, closer study of the first 24 hour period revealed distinct early colonization defects in each mutant (Figure 7B). The Δ*bcsC* mutant was recovered at lower levels at the earliest time points but reached comparable densities to wild-type by 24 hours (Figure 7B). Genetic complementation of Δ*bcsC* by expressing the *bcs* operon from an ectopic locus resulted in a colonization phenotype that was indistinguishable from wild-type at all time points. We thus conclude that cellulose contributes to initial RL271 root colonization but is dispensable for its maintenance on root tissue. The Δ*fcl* capsule mutant colonized like wild-type initially, but its population expanded more slowly, resulting in reduced recovery at 16 and 24 hours (Figure 7B); populations reached wild-type levels at later time points (Figure 7A) consistent with a transient colonization defect. Wild-type colonization kinetics in Δ*fcl* were restored by ectopic expression of *fcl*. The Δ*fcl* Δ*bcsC* double mutant exhibited the combined defects of both single mutants, with early colonization defects similar to Δ*bcsC* and slower population expansion similar to Δ*fcl*. Together, these results demonstrate that capsule and cellulose each support root colonization by RL271, specifically during early and intermediate stages of establishment. However, neither polysaccharide is strictly required for long-term root association in this sterile plant model.

**Figure 7:**
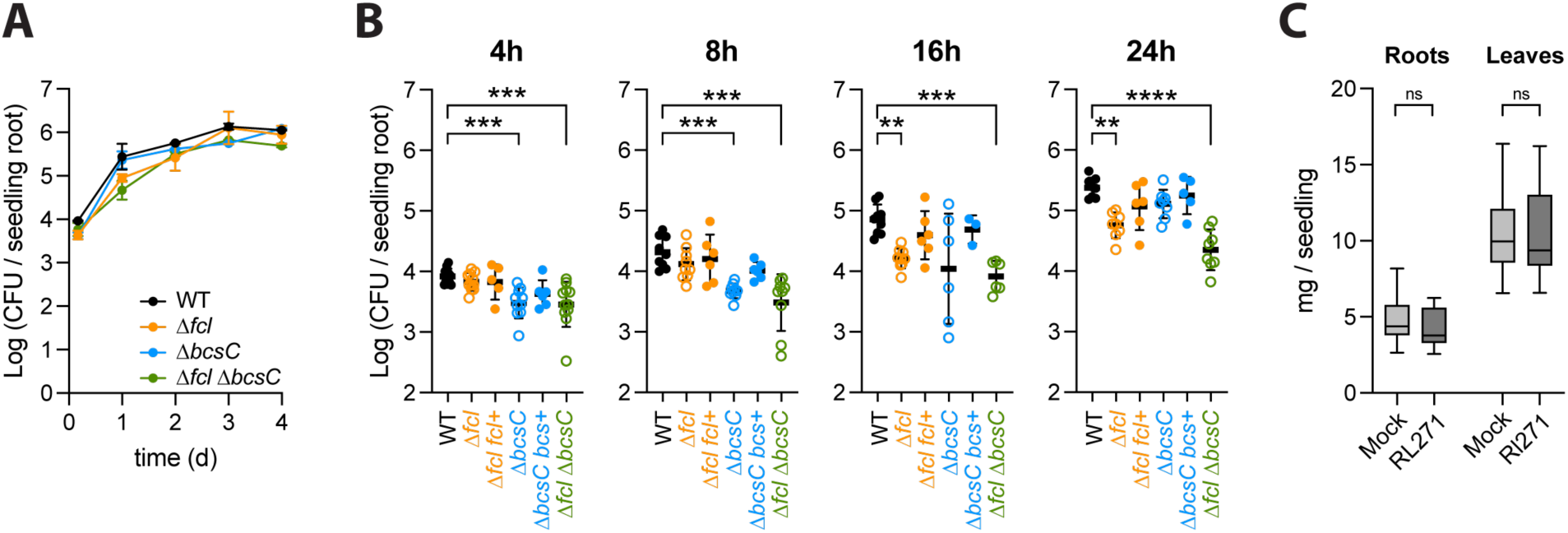
*Caulobacter* sp. RL271 efficiently colonizes *Arabidopsis* roots and cellulose synthesis promotes early colonization. Colonization of sterile 1-week old *Arabidopsis* seedlings inoculated with wild type or mutant *Caulobacter* sp. RL271. **(A)** CFU enumerated from the roots of seedlings inoculated with wild type, Δ*fcl*, Δ*bcsC* or Δ*fcl* Δ*bcsC* strains, harvested 4 hours, 1 day, 2 days, 3 days, or 4 days post-inoculation. Each time point reflects the mean ± SD of the seedlings on two replicate plates (5 seedlings per plate). **(B)** CFU enumerated from the roots of seedlings inoculated with wild type, Δ*fcl*, Δ*fcl glmS*::*fcl* (Δ*fcl fcl*+), Δ*bcsC*, Δ*bcsC glmS*::*bcs operon* (Δ*bcsC bcs*+), or Δ*fcl* Δ*bcsC* strains, harvested, 4, 8, 16, or 24 hours post-inoculation. Each point reflects the mean CFU per seedling of the 5 seedlings grown together on a plate. The black bars represent the mean ± SD of 6 to 11 replicate plates (5 seedlings per plate) collected over 3 independent experiments. Within each time point, strains were compared to wild-type with a Kruskal-Wallis test followed by a Dunn’s multiple comparisons test. Strains that were significantly different from wild-type are indicated: ** p <0.01; *** p < 0.001; **** p< 0.0001. **(C)** Biomass of roots (left) or leaves and stems (right) of seedlings inoculated with wild type RL271 (dark grey) or mock inoculated with sterile broth (light grey) one week post inoculation. Each box plot represents mass of seedlings from 18-20 plates per group (5 seedlings per plate) collected over three independent experiments. Whiskers indicate min and max. For each tissue type, inoculated and mock-inoculated masses were compared with two-tailed unpaired t-test; ns = p > 0.05.

### Caulobacter sp. RL271 colonizes Arabidopsis thaliana roots without directly promoting plant growth

Having established that RL271 colonizes *Arabidopsis* roots, we next asked whether colonization has a direct effect on plant growth. This question was motivated in part by our prior observations that inoculation of soil with RL271 enhanced soybean growth during drought (Longley, 2022), raising the possibility that RL271 has direct plant growth-promoting properties. To test this, we conducted an assay similar to that used by Luo and colleagues, in which *Caulobacter* sp. RHG1, a maize root isolate, was shown to promote *Arabidopsis* seedling growth under sterile conditions (Luo et al., 2019). Seven-day-old *Arabidopsis* seedlings were inoculated with RL271 or mock-inoculated with PYE growth medium, and shoot and root biomass were measured 7 days post-inoculation. We detected no significant difference in biomass between inoculated and uninoculated plants (Figure 7C).

## Discussion

*Caulobacter* sp. RL271 represents a distinct terrestrial *Caulobacter* lineage whose biology is best understood in the context of plant-associated *Caulobacter* spp. rather than comparison to the classical aquatic model *C. crescentus.* Its large genome, higher GC content, and phylogenetic placement near the root isolate CBR1 are consistent with the broader genomic and ecological distinction between terrestrial and aquatic *Caulobacter* lineages (Wilhelm, 2018; Hallgren et al., 2025). With the closest genome being *Caulobacter* sp. CBR1 having 91.8% ANI, and with ∼86% ANI to *C. segnis* and *C. crescentus*, RL271 and CBR1 appear to represent a distinct, unnamed lineage that has retained cellular dimorphism but diverged substantially in nutritional and surface-associated traits.

### Loss of holdfast and gain of a capsule-cellulose matrix define the surface biology of RL271

Several traits of RL271 echo the *C. segnis* type strain, including reduced stalk formation, absence of the riboflavin biosynthesis operon, and absence of holdfast genes (Urakami et al., 1990; Patel et al., 2015). The co-occurrence of these features in RL271, CBR1, and *C. segnis* suggests they reflect shared selective pressures in plant-associated or soil environments, and points toward a coherent ecological syndrome within this clade. Recent phylogenomic work has shown that lifecycle complexity and polar surface traits are evolutionarily labile across the Caulobacterales, with repeated reductions and losses of classical dimorphic features among relatives of *C. crescentus* (Hallgren et al., 2025). RL271 extends this picture in an important way. Specifically, loss of holdfast does not imply loss of a structured surface associated lifestyle. Instead, holdfast can be functionally replaced by a matrix architecture built from capsule and cellulose that similarly serve to mediate surface interactions in a different habitat.

The significance of holdfast loss extends beyond cell biology. Holdfast-associated genes have been considered “signature genes” of the order *Caulobacterales* and used as diagnostic markers for *Caulobacter* in metagenomic surveys (Gupta and Mok, 2007; Wilhelm, 2018), and holdfast-mediated adhesion is central to how surface colonization has been conceptualized in *C. crescentus*, where the holdfast provides effectively irreversible attachment to solid substrates (Merker and Smit, 1988; Tsang et al., 2006; Berne et al., 2013). Moreover, enrichment in a film at the air-liquid interface has historically been used to facilitate isolation of *Caulobacter* (Poindexter, 1964, 1981), and this process relies on holdfast (Fiebig, 2019). Holdfast-deficient lineages like RL271 and CBR1 are therefore likely undercounted in surveys that rely on these markers, meaning the true prevalence of *Caulobacter* in plant-associated environments may be higher than current estimates suggest. The selective advantage of losing holdfast is not immediately obvious, but there is relevant precedent in laboratory evolution. The highly studied strain *C. crescentus* NA1000 carries a spontaneous frameshift in *hfsA* that abolishes holdfast production; this mutation became fixed in the population over years of serial passage in the laboratory because non-adherent cells are passaged more efficiently under routine culture conditions (Marks et al., 2010). Deliberate adhesion selection experiments can similarly enrich a spectrum of holdfast-deficient mutants (Ong et al., 1990; Hershey et al., 2019a) These observations establish that holdfast production carries a measurable fitness cost when permanent attachment provides no advantage (or is disadvantageous). In the root endosphere, where bacteria must navigate intercellular spaces rather than anchor permanently to exposed surfaces, analogous selection against holdfast production is plausible.

In place of holdfast, RL271 deploys cellulose and capsular polysaccharide in functionally opposing but complementary roles. Cellulose drives aggregation and provides structural cohesion, while capsule limits close cell-cell contact and maintains an open biofilm architecture. This differs fundamentally from the *C. crescentus* paradigm, where a localized polar adhesive mediates permanent surface attachment, but it is consistent with growing recognition that extracellular polymers tune not only adhesion but also the spatial mechanics of bacterial communities (Dragos and Kovacs, 2017). RL271 exhibits the opposite adhesive balance from *C. crescentus*. It has strong cell-cell cohesion but little persistent attachment to glass or plastic. This adaptation may be less for permanent anchoring to abiotic surfaces than for persistence in hydrated, particle-rich, or host-associated microenvironments where reversible aggregation and community integration are advantageous. The strong similarity between the RL271 capsule biosynthesis locus and the capsule-related mobile genetic element of *C. crescentus* NA1000, together with the presence of a *hvyA* ortholog (Ardissone et al., 2014) nearby on the chromosome, suggests that RL271 has retained a conserved capsule module but deployed it in a distinct ecological context. Whether the RL271 capsule also confers phage resistance, as it does in NA1000, is an open question of particular interest. The apparent essentiality of multiple capsule-associated genes in the transposon dataset is consistent with prior observations that partial disruption of polysaccharide biosynthesis pathways can generate toxic intermediates or envelope stress rather than reflecting a true absolute growth requirement (Ardissone et al., 2014).

The root colonization phenotypes of Δ*bcsC* and Δ*fcl* reinforce the roles of these matrix components during early establishment. Root association is a multistep process in which extracellular polymers often contribute most critically to the transition from initial surface contact to stable microcolony formation (Knights et al., 2021). In RL271, neither cellulose nor capsule was absolutely required for long-term root association, but cellulose and capsule mutant strains had diminished colonization kinetics, consistent with roles in early attachment and community integration rather than in maintenance of an established population. The cellulose phenotype in particular aligns with roles described in other plant-associated bacteria, where cellulose strengthens attachment and biofilm formation on root surfaces (Matthysse et al., 2005). This role in early attachment may be crucial in complex soil microbiomes, allowing early establishment and niche colonization prior to the arrival of other taxa.

### Riboflavin auxotrophy, cross-feeding, and the Black Queen hypothesis

The genomic loss of the riboflavin biosynthesis operon in RL271 has a clear ecological consequence. RL271 requires exogenous riboflavin for growth, and proximity to *B. diazoefficiens* is sufficient to rescue this requirement under riboflavin-limiting conditions, providing evidence that *Bradyrhizobium*-derived flavins can support RL271 in the shared root environment. This finding provides a concrete mechanistic example of bacteria-bacteria facilitation in a root-associated *Caulobacter*. It fits well within a broader literature showing that rhizobia constitutively secrete riboflavin and related flavins into the rhizosphere (Phillips et al., 1999; Dakora et al., 2015), that the level of flavin secretion influences rhizobial root colonization efficiency (Yurgel et al., 2014), and that vitamin-producing taxa can shape rhizosphere community assembly by supporting auxotrophic members that could not otherwise persist (Garrido-Sanz and Keel, 2025). The requirement for capsule in the directed growth of RL271 toward *Bradyrhizobium* on riboflavin-limiting surfaces suggests that RL271 may actively position itself relative to its riboflavin source rather than simply benefiting from passive diffusion. This observation connects the surface matrix biology of RL271 directly to its nutritional ecology, and raises the possibility that capsule-mediated spreading serves a foraging function in riboflavin-limited environments. Together, these results are compatible with Black Queen dynamics, under which the reliable leakage of a costly metabolite into the shared environment can favor adaptive loss of the corresponding biosynthetic function in community members that can exploit it (Morris et al., 2012).

RL271 satisfies two parts of this model: *a)* it has lost riboflavin biosynthesis and *b)* it can exploit extracellular flavin generated by *Bradyrhizobium*. The sporadic distribution of riboflavin operon loss across *Caulobacter* lineages (Hallgren et al., 2025) is consistent with repeated independent gene loss events and suggests that reliable exogenous flavin sources are available across a range of *Caulobacter* habitats. This pattern is what the Black Queen hypothesis predicts. However, whether operon loss in each case was actively selected for because community-level flavin availability was sufficient to compensate, or whether the operon decayed neutrally and the resulting dependency subsequently shaped ecological associations, cannot yet be distinguished. We interpret our RL271 data as evidence for a flavin-dependent interspecies dependency consistent with a Black Queen-like scenario, while acknowledging that the evolutionary drivers of operon loss in this lineage remain to be established.

### On the hub taxon function of RL271

RL271 colonized *Arabidopsis* roots without measurably increasing seedling biomass under sterile conditions. This contrasts with *Caulobacter* sp. RHG1, a maize root isolate that promotes *Arabidopsis* growth in an analogous sterile system (Luo et al., 2019), and is an important reminder that root colonization and plant growth promotion are distinct phenomena that should not be conflated. The mechanisms by which RHG1 promotes plant growth remain incompletely characterized, but its ability to do so in a host-independent, microbiome-independent context indicates that it acts through a direct plant-signaling or nutrient-provisioning mechanism. That RL271 lacks this capacity on *Arabidopsis* seedlings suggests that its beneficial effects on soybean performance, including enhanced nodule activity, elevated *Bradyrhizobium* levels, and increased biomass under water limitation (Longley, 2022), depend on host identity, environmental context, stress conditions, or interactions with *Bradyrhizobium* and other microbiome members rather than a more broad plant growth-promoting ability. This distinction is consequential. It implies that RL271 functions as an ecological facilitator rather than a direct growth promoter, with its hub-taxon status arising from its position at the interface of a syntrophic bacterium-bacterium interaction.

Perhaps the most interesting unresolved question is the nature of the relationship between RL271 and *Bradyrhizobium*. Data presented here provide evidence that *Bradyrhizobium* supplies riboflavin to RL271, and a companion study demonstrates that RL271 in turn supports *Bradyrhizobium* growth through a mechanism that remains to be fully characterized (Longley, 2022). Together, these results reframe the *Caulobacter* sp. RL271-*Bradyrhizobium* interaction as a genuine mutualism rather than a one-sided dependency (commensalism), and raise broader questions about how such pairwise microbial interactions scale to influence plant performance at the community level. RL271 alone was sufficient to recapitulate the growth-promoting effects of a five-taxon hub consortium in soybean (Longley, 2022) indicating that its ecological influence is disproportionate to its numerical abundance, a hallmark of keystone taxa in microbiome networks. Understanding the molecular basis of the RL271-*Bradyrhizobium* mutualism, through transcriptomic profiling of both partners during root colonization, defined co-inoculation experiments in soybean, and targeted genetic dissection of candidate interaction mechanisms, will be essential for understanding how hub taxa shape the productivity of legume-microbiome systems and can inform principled strategies to leverage microbial interactions in sustainable agriculture.

## Materials and Methods

### Growth conditions

*Escherichia coli* strains were grown at 37°C with lysogeny broth (LB) [10 g/L peptone, 5 g/L yeast extract, 5 g/L NaCl]. *Caulobacter* strains were grown at 30°C using either PYE medium [2 g/L peptone, 1 g/L yeast extract, 1 mM MgSO4, 0.5 mM CaCl2], R2A medium (premix from Teknova) (Reasoner and Geldreich, 1985), or soil extract medium [20% soil extract, 1 g/L yeast extract, 5 g/L xylose or arabinose). *Bradyrhizobium* was grown at 30°C using either R2A or soil extract medium. Soil extract was prepared according to the DSMX recipe 98 (*Rhizobium* medium), except that soil was not dried prior to use. Briefly, approximately 400 g damp soil and 1 g Na2CO3 were mixed with 1 L water and autoclaved for 60 minutes at 121°C and >15 PSI. After the soil particulates settled overnight, the broth was decanted from the sediment and then centrifuged for 15 minutes at 10,000 rcf to remove fine particulates. The extract was then decanted from the pelleted particulates and stored in 50 ml aliquots at −20°C until use. As necessary, growth media were solidified with 15 g/L agar, and/or supplemented with antibiotics or DAP. Unless otherwise noted, growth media for *Caulobacter* sp. RL271 were either freshly prepared or supplemented with 1-5 uM riboflavin. Antibiotic concentrations were as follows: Kanamycin 50 µg/ml (*E. coli*), 25 µg/ml (*Caulobacter* solid media), 5 ug/ml (*Caulobacter* liquid media), Carbenicillin 100 µg/ml (*E. coli*). As needed, DAP was added at a final concentration of 300 µM. Unless noted otherwise, cultures were grown in 2 ml of growth medium in 14 X 100 mm glass culture tubes at a 45° angle shaking at 200 rpm.

### Plasmid construction and strain building

Plasmids were constructed using standard molecular biology techniques briefly described here. Full descriptions of all plasmids and strains used in this study can be found in Table S1. Desired genomic regions were amplified from RL271 DNA using KOD Xtreme HotStart Polymerase (Novagen) following manufactures instructions and primers with appended restrictions sites. When necessary, DNA fragments were joined by overlap-extension PCR (Horton et al., 1989). PCR products were digested with appropriate restriction enzymes (NEB) and ligated into similarly digested plasmids with T4 DNA ligase (NEB). Ligated products were transformed into chemically competent *E. coli* TOP10 and transformed clones were selected on LB containing the appropriate selective antibiotic. The resulting plasmids were confirmed via Nanopore sequencing (Plasmidsaurus), and transformed into *E. coli* WM3064, a conjugation competent, DAP auxotroph.

To engineer deletions in the RL271 chromosome, we used two-step recombination with sucrose counterselection following the same approach as is widely used for *C. crescentus*. Approximately 500 bp upstream and downstream of the region to be deleted were cloned into pNPTS138. The resulting plasmids were conjugated from WM3064 into *Caulobacter sp.* RL271. Primary integrants were selected on PYE agar containing 25 µM kanamycin and 1 µM riboflavin. Individual KanR colonies were grown non-selectively in PYE broth with 1 µM riboflavin for 6-18 hours and then plated on PYE containing 3% sucrose and 1 µM riboflavin to select for secondary recombinants. Sucrose resistant colonies were patched on media with and without kanamycin to confirm kanamycin sensitivity and then screened by PCR to distinguish secondary recombinants in which the wild-type allele was deleted from those in which it was restored.

To complement deleted genes, we used a miniTN7 transposon carrying a kanamycin resistance cassette. Genes were PCR amplified with ∼300 bp upstream to capture their native promoter and cloned into the TN7 cassette of the plasmid pUC18-miniTN7k (Myeni et al., 2013). The resulting plasmids were co-conjugated from WM3064 into RL271 with pTNS3, a helper plasmid encoding the TN7 transposase (Choi et al., 2008). Strains in which the TN7 cassette integrated in the RL271 genome were selected on PYE agar containing 25 µM kanamycin.

### *Caulobacter* sp. RL271 genome sequencing and assembly

The complete *Caulobacter* sp. RL271 genome was generated by a hybrid assembly of short- and long-read sequences. Residual adapter sequences were trimmed from Oxford Nanopore Technology (ONT) reads using Porechop v0.2.4 (https://github.com/rrwick/porechop). De novo genome assembly was performed from the trimmed ONT reads using Flye v2.9.2 (Kolmogorov et al., 2019) under the nano-hq model, with assembly coverage set to 50x and an estimated genome size of 6 Mb. Subsequent polishing used paired-end Illumina short reads aligned with Bowtie2 v2.5.1 and polished with Pilon v1.24 (Walker et al., 2014) under default parameters. Long-read contigs with an average short-read coverage of 15x or less were removed to reduce assembly artifacts attributable to low-quality nanopore reads. Assembled contigs were evaluated for circularization using Circulator v1.5.5 (Hunt et al., 2015) with ONT long reads. This approach yielded a single circular chromosome of 5,687,505 bp with no detected plasmids. The complete genome sequence is available through NCBI GenBank accession CP096040. Reads used to assemble the genome are available at the NCBI BioProject Accession PRJNA828140.

### *Caulobacter* spp. phylogenetics

We analyzed 34 *Caulobacter* genomes and 1 *Brevundimonas* genome as an outgroup (see Table S2 for genome accession numbers and select metadata on each genome). Single-copy, conserved protein-coding markers were identified with GToTree v1.8.16 (Lee, 2019) using the built-in Alphaproteobacteria HMM set (117 targets). Default GToTree settings were used. GToTree extracted marker proteins from each genome, aligned each locus, and produced a concatenated amino-acid supermatrix plus a per-gene partition file. The final concatenated alignment contained 24,579 amino-acid sites across the 35 taxa. Phylogenetic inference was performed with IQ-TREE2 v2.2.2.7 (Minh et al., 2020) using the alignment matrix (117 partitions). Substitution models were selected with ModelFinder (Kalyaanamoorthy et al., 2017) using MFP+MERGE. Node support was assessed with ultrafast bootstrap (UFBoot2) (Hoang et al., 2018) using 1,000 replicates and Nearest Neighbor Interchange (NNI) correction.

### Construction of a barcoded himar transposon mutant library

A barcoded transposon library was constructed in *Caulobacter* sp. RL271 by conjugation with the *E. coli* donor strain APA752, which carries the pKMW3 Himar transposon vector library (gift from Adam Deutschbauer, University of California Berkeley). A 2 ml starter culture of RL271 was inoculated into 20 ml of 2X PYE supplemented with 1 µM riboflavin, and outgrown for approximately 6 hours at 30°C with shaking. In parallel, one 1 ml aliquot of an APA752 freezer stock was inoculated into 20 ml LB containing 50 µg/ml kanamycin and 300 µM DAP for 6 hours at 37°C with shaking. The two cultures were combined, pelleted by centrifugation (7,000 x g, 5 min), resuspended in 100 µl fresh PYE, and spotted onto a PYE agar plate supplemented with 300 µM DAP and 1 µM riboflavin. After overnight incubation at 30°C, cells were scraped from the plate, resuspended in 1.5 to 2 ml PYE, and spread across 16 large (150 mm) PYE agar plates containing 1 µM riboflavin and 5 µg/ml kanamycin (100 µl of bacterial suspension per plate) to select for RL271 transconjugants. After 4 days at 30°C, an estimated 10⁵ colonies were scraped from these plates and resuspended in PYE broth. The pooled cells were used to inoculate 300 ml of 2X PYE containing 1 µM riboflavin and 25 µg/ml kanamycin to a starting density of OD660 = 0.1 and outgrown for 9 hours to OD660 = 0.6. Cells were pelleted by centrifugation (7,000 x g for 5 min), resuspended in 30 ml PYE containing 20% glycerol, and frozen in aliquots at −70°C. Two 1 ml aliquots were reserved for genomic DNA extraction to map transposon insertion sites.

### Mapping Tn-Himar insertion sites in the *Caulobacter* sp. RL271 mutant library

Transposon insertion sites were mapped following published protocols (Wetmore et al., 2015) with modifications to the PCR enrichment strategy outlined below. Genomic DNA was extracted from the mutant pool and about 10 µg was sheared to approximately 300 bp fragments in a volume of 130 µl using a Covaris M220 ultrasonicator under manufacturer’s settings. Sheared DNA (1.25 µg) was end-repaired, A-tailed, and ligated to a custom Y-adapter prepared by annealing oligonucleotides Mod2_TruSeq and Mod2_TS_Univ (Table S1), using the NEBNext Ultra II Library Prep Kit (New England Biolabs, E7103S) per the manufacturer’s protocol. The ligated product was cleaned by two-sided SPRIselect bead selection (0.5-0.3X) and eluted in 35 µl of 10 mM Tris, pH 8.5.

Fragments containing Himar transposons were enriched by two-step nested PCR using GoTaq Green Master Mix (Promega, M7122). Primer sequences are listed in Table S1. The first reaction used the forward primer TS_phimar+4, which is derived from TS_phimar (Wetmore et al., 2015). This primer incorporates the Illumina TruSeq Read 1 sequence, a random hexamer to facilitate clustering, and an extended transposon-specific sequence for improved specificity. The reverse primer, TS_R, containing the TruSeq Read 2 sequence complementary to the adapter. Each reaction contained 0.3 µM of each primer, 15 µl of adapter-ligated template, 5% DMSO, and 1X GoTaq in 100 µl total volume, with the following cycling conditions: 98°C for 3 min; 20 cycles of 98°C for 30 s, 65°C for 20 s, and 72°C for 30 s; 72°C for 10 min. The product was cleaned with 0.85X SPRIselect beads and eluted in 40 µl of 10 mM Tris, pH 8.5. The second reaction appended Illumina P5 and P7 sequences and introduced a 6-bp index on the P7 end, using primers P5_TS_F and P7_MOD_TS_index8 (0.5 µM each), 5 µl of the first-reaction product, 5% DMSO, and 1X GoTaq in 100 µl, with the following cycling conditions: 98°C for 3 min; 15 cycles of 98°C for 20 s, 69°C for 10 s, and 72°C for 20 s; 72°C for 5 min. The final product was cleaned with 1X SPRIselect beads and sequenced on an Illumina NovaSeq X Plus.

Sequences were analyzed using custom scripts (Wetmore et al., 2015) available at https://bitbucket.org/berkeleylab/feba/src/master/. Insertion sites were aligned and mapped to the RL271 genome using BLAT (Kent, 2002), and unique barcode sequences were linked to their corresponding genomic locations using MapTnSeq.pl. Barcodes mapping consistently to a single genomic location were identified using DesignRandomPool.pl. Raw sequence Tn-seq reads are deposited in the NCBI Sequence Read Archive under BioProject accession PRJNA828140.

### Visual screen of transposon mutants

An aliquot of the TN-himar mutant pool was thawed, outgrown in PYE for several hours, serially diluted and spread on 150 mm petri plates containing solidified PYE or R2A at a density of approximately 500-1000 colonies per plate. After 2-3 days of growth at 30°C, colonies were visually screened to identify mutants that were more or less mucoid than wild type. We note that mutants with changes in colony appearance were more obvious on R2A than PYE. Candidate mutants were regrown on R2A agar to confirm aberrant colony morphology. Transposon insertion sites were mapped in mutants with confirmed colony morphology defects.

### Mapping transposon insertion sites in individual mutant clones

We used arbitrary-nested PCR, combined with sanger sequencing to identify the transposon insertion junctions with the chromosome, adapting a previously described approach (O’Toole et al., 1999). Briefly, a first round PCR reaction contained a primer complementary to the transposon (U1-fw) and an arbitrary primer with a random heptamer followed by a standard sequencing primer (M13-N7) for priming from the chromosome (see Table S1 for primer sequences). The template consisted of a small colony of cells resuspended in 50 ul of water. Each reaction contained 0.3 µM of each primer, 1 ul resuspended cells, 1X GoTaq (Promega) in a total volume of 20 µl, with the following cycling conditions: 95°C for 2 min; 35 cycles of 95°C for 30 s, 38°C for 30 s, and 72°C for 60 s; 72°C for 5 min. The primary PCR product is often not visible on a gel. Primary PCR products were treated with ExoSAP-IT (Applied Biosystems) to remove excess primer. The second round PCR reaction contained a transposon specific primer 3’ of U1-fw priming site (U2-out) and the M13F primer, which will anneal to fragments amplified with the M13-N7 primer in the first round. Each reaction contained 0.3 µM of each primer, 2 ul ExoSAP treated primary PCR product and 1X GoTaq in total volume of 20 µl, with the following touchdown cycling conditions: 95°C for 2 min; 35 cycles of 95°C for 30 s, 68-48°C (starting at 68°C and decreasing by 0.5°C each cycle) for 30 s, and 72°C for 60 s; 72°C for 5 min. PCR products were again cleaned enzymatically with ExoSAP-IT, and then Sanger sequenced using the U2-out primer. Insertion sites were identified by comparing the 30 bp downstream of the end of the transposon (CAACCTGT**TA**) to the RL271 genome by BLAST (Johnson et al., 2008).

### Growth curves

To monitor growth kinetics, cultures at a starting density of 0.01 OD660 were transferred to 24- or 48-well microtiter plates. A transparent lid was sealed on the plate with strips of AeraSeal (Excel Scientific) to prevent evaporation. In a Tecan Infinite M Nano plate reader, cultures were incubated at a 30 C with shaking and the optical density at 660 nm was measured every 10 minutes.

### Crystal violet staining of surface attached cells

To assess formation of surface attached biofilms, cells were inoculated into 1 ml of PYE broth in 24 well plates at a starting optical density of 0.01 OD660. Plates were sealed with AeraSeal (Excel Scientific) and grown either with shaking (150 rpm) or without shaking overnight at 30°C. Plates were photographed to document culture growth, then cells were removed by aspiration. Wells were gently washed with a stream of water, filled with 1.5 ml of 0.01 % crystal violet, and incubated at RT for 5 min. The dye was removed by aspiration and the wells were rinsed with water before photographing the wells again.

### Cell staining and widefield light microscopy

Cells were imaged as wet mounts at 630 X magnification using a Leica DMI 6000 microscope with an HC PL APO 63× / 1.4 numeric aperture oil Ph3 CS2 objective. Images were captured with an Orca-ER digital camera (Hamamatsu) controlled by Leica Application Suite X (Leica). Fluorescence images were captured using the filter cubes indicated below. Images were scaled and cropped with either Leica Application Suite X or Fiji (Schindelin et al., 2012).

#### Wheat-germ agglutinin staining

To stain holdfast polysaccharide, cells were grown to log phase (approximately 0.2 OD660) in PYE. 500 ul of cells were mixed with Wheat Germ Agglutinin Alexa Fluor 594 conjugate (WGA-Alexa594) (Invitrogen W1262) to a final concentration of 5 ug/ml and incubated at room temperature in the dark for 5-10 minutes. Cells and excess dye were diluted with 1 ml PYE and then centrifuged for 2 min at 10,000 x *g* to concentrate cells. After removing the supernatant, the cells were resuspended in the remaining 20-30 ul of broth. WGA-Alexa594 fluorescence was visualized using a Leica TX2 filter cube (Excitation: BP 560/40; Dichroic: 595; Emission: BP 645/75).

#### Dextran exclusion

FITC-dextran with average mass of 2000 kDa (Sigma-Aldrich FD2000S) was used to assess capsule thickness as previously described (Ardissone et al., 2014). Briefly, cells were grown in PYE to log phase (approximately 0.2 OD660). The cells in 500 μl of each culture were harvested by centrifugation at room temperature (3000×*g*, 5 min), washed once with PBS, and resuspended in 30 μl of PBS. Then 10 μl of bacterial suspension was mixed with 2 μl of FITC-dextran (10 mg/ml in water). One µl was applied onto a microscope slide and firmly covered with a coverslip. Fluorescence was visualized using L5 filter cube (Excitation: BP 480/40; Dichroic: 505; Emission: BP 527/30). We used Fiji (Schindelin et al., 2012) to identify and measure cell areas in phase contrast images and dextran excluded areas in fluorescence images.

#### Calcofluor white staining

To visualize cellulose, cells were scraped from PYE agar plates after 2 days of growth at 30°C, resuspended in 90µl liquid PYE, and mixed with 10µl 0.01% calcofluor white (CW) solution to achieve a final concentration of 0.001% CW. The suspension was incubated at room temperature for 15 minutes, then diluted with 900µl water to serve as a wash step. Cells were pelleted by centrifugation at room temperature (6000×*g,* 3 minutes), the supernatant was removed, and the cells were resuspended in the 20-30 μl of remaining liquid. CW fluorescence was visualized using a Leica DAPI filter cube (Excitation: BP 350/50; Dichroic: 400; Emission: BP 460/50).

### Scanning electron microcopy

Starter cultures were inoculated into fresh PYE supplemented with 1 µM riboflavin at a starting density of 0.01 OD660. One ml of diluted culture was transferred into wells of 24-well plates containing a sterile 12-mm round coverslips. Cultures were grown overnight at room temperature without shaking. After growth, 1 ml of 4% glutaraldehyde was gently added to each well, followed by a 30 minute incubation to allow sample fixation. The coverslips were carefully removed to minimally disrupt associated cells and placed in a graded ethanol series (25%, 50%, 75%, 95%) for ten min at each step and with three 10 min changes in 100% ethanol (Klomparens et al., 1986). Samples were critical point dried in a Leica Microsystems model EM CPD300 critical point dryer (Leica Microsystems, Vienna, Austria) using carbon dioxide as the transitional fluid. Coverslips were mounted on aluminum stubs using System Three Quick Cure 5 epoxy glue (System Three Resins, Inc., Aubur, WA). Samples were coated with osmium (≈10 nm thickness) in a Tennant20 osmium CVD (chemical vapor deposition) coater (Meiwafosis Co., Ltd., Osaka, Japan) and examined in a JEOL 7500F (field emission emitter) scanning electron microscope (JEOL Ltd., Tokyo, Japan) at the Michigan State University Center for Advanced Microscopy.

### Plant growth and inoculation

Overnight cultures of Caulobacter sp. RL271 grown in PYE broth were normalized to a density of 0.01 OD660 (approximately 5 X 10^6^ CFU/ml) in sterile PYE. Prior to inoculation, axenic *Arabidopsis thaliana* accession *Col-0* seeds were surface-sterilized with 70% ethanol and 5% bleach (1 min each), followed by thorough washes with sterile water. Sterile seeds were stratified at 4°C in the dark for 3 days to synchronize germination. Seedlings were then sown onto 0.5X Murashige and Skoog medium with Gamborg’s B5 vitamins (MS) (Sigma) and 30 g/L sucrose solidified with 6 g/L Phyto Agar (GoldBio) and placed in a growth chamber (22°C, 10h light/18°C, 14h dark) for 7 days to germinate. Following germination, five seedlings were transferred to square plates (100mm x 100mm) containing 0.25X MS solidified with 10 g/L Phyto Agar for vertical growth in the same light and temperature regime. Before transferring the seedlings, the vertical growth plates were inoculated with 100µl of bacterial culture (approximately 5 X 10^5^ CFU), or a sterile PYE control, evenly spread across the surface.

#### Plant biomass measurements

After transferring to vertical plates as described above, seedlings were grown as described for 7 days. At the time of harvest, shoots and roots were separated at the hypocotyl, and tissues were immediately placed tissue into pre-weighed 1.5ml Eppendorf tubes. The five seedling shoots or 5 seedling roots from each plate were combined in one tube. The tubes were then re-weighed, and the difference was calculated to determine fresh tissue mass. The mass was divided by five to calculate the average fresh mass per seedling for shoots and roots.

#### Plant colonization assays

Seedlings were grown vertically as described, and samples were harvested at 4, 8, 16, or 24 hours (early colonization) or 1, 2, 3, or 4 days (late colonization). At the time of harvest, roots were sectioned at the hypocotyl, pooled per plate, and transferred to 1.5ml screw cap red RINO tubes (Next Advance) containing 500µl of sterile PYE. Root tissue was homogenized using a Bullet Blender Storm Pro Homogenizer (Next Advance) for 10 minutes at speed 10. Homogenates were 10-fold serially diluted (10^-1^ to 10^-8^) in sterile PYE. Each dilution was plated in triplicate 5µl spots on PYE agar. CFUs per seedling root were enumerated after 1-2 days of incubation at 30°C.

## Supporting information

Supplemental Table 1: Strains, plasmids and oligos

Supplemental Table 2: Caulobacter genome accessions and metadata

RL271 essential gene lists

## Acknowledgements

We thank Amy Albin in the MSU Center for Advanced Microscopy for assistance with scanning EM, and Miette Hennessy for assistance with the visual screen.

## Author contributions

S.C. G.B. R.L. and A.F. conceived the study; M.A.Q., S.C. and A.F designed and performed the experiments and analysis; S.L.L assisted with methodology; M.A.Q., S.C. and A.F wrote the original draft; M.A.Q, S.C., S.L.L, G.B, R.L, and A.F. reviewed and edited the manuscript. All authors approved the final manuscript.

## Supplemental Materials

Table S1: Strains, plasmids and oligos used in this study

Table S2: *Caulobacter* genome accessions and meta data corresponding to isolates used in Figure 1

Table S3: *Caulobacter sp* RL271 essential gene predictions from TnSeq

## Supplemental Figures

**Figure S1:**
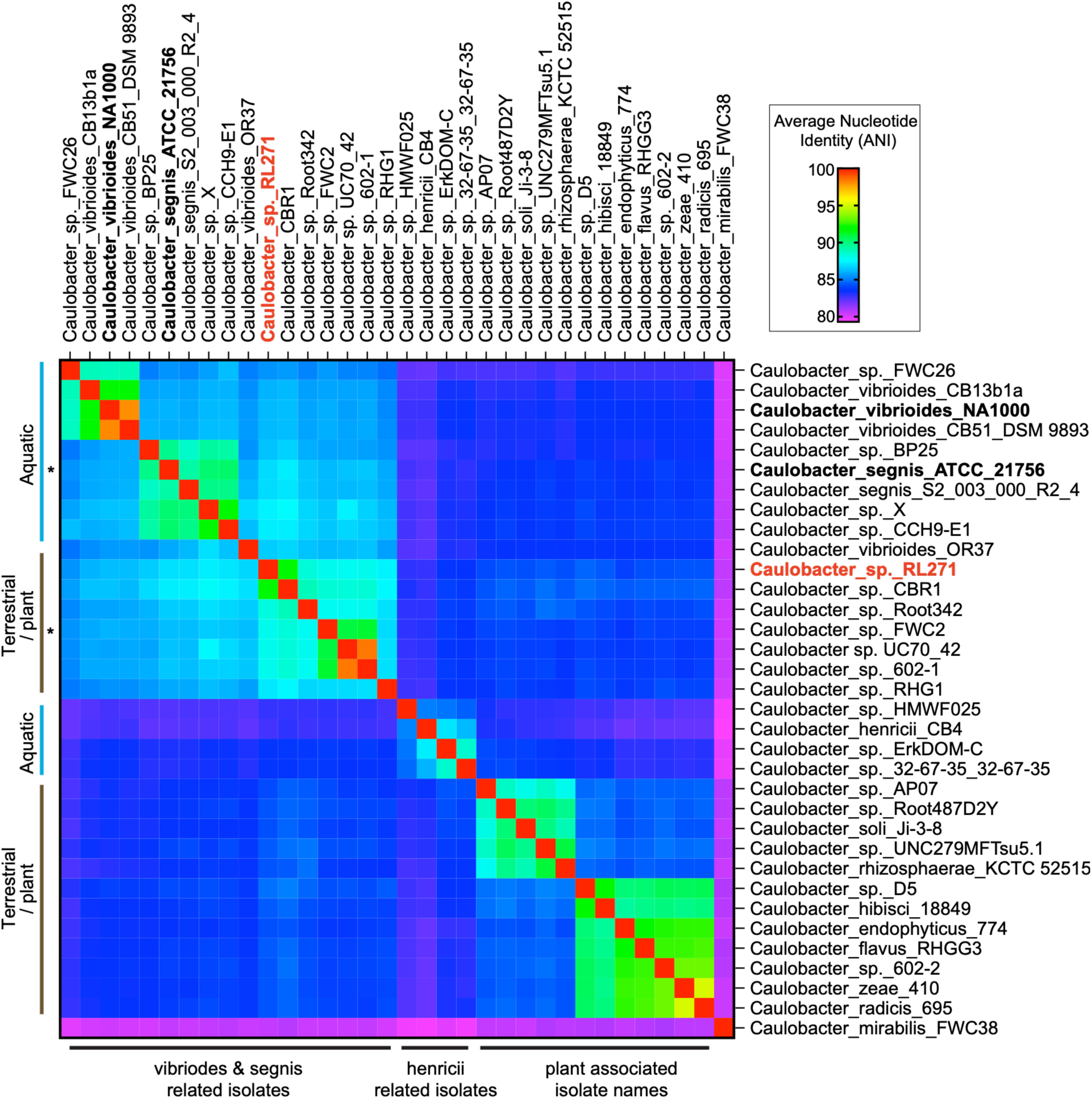
Heatmap of pairwise average nucleotide identity between *Caulobacter* sp. genomes in Figure 1. Genome accession information for each isolate is in Table S2.

**Figure S2.**
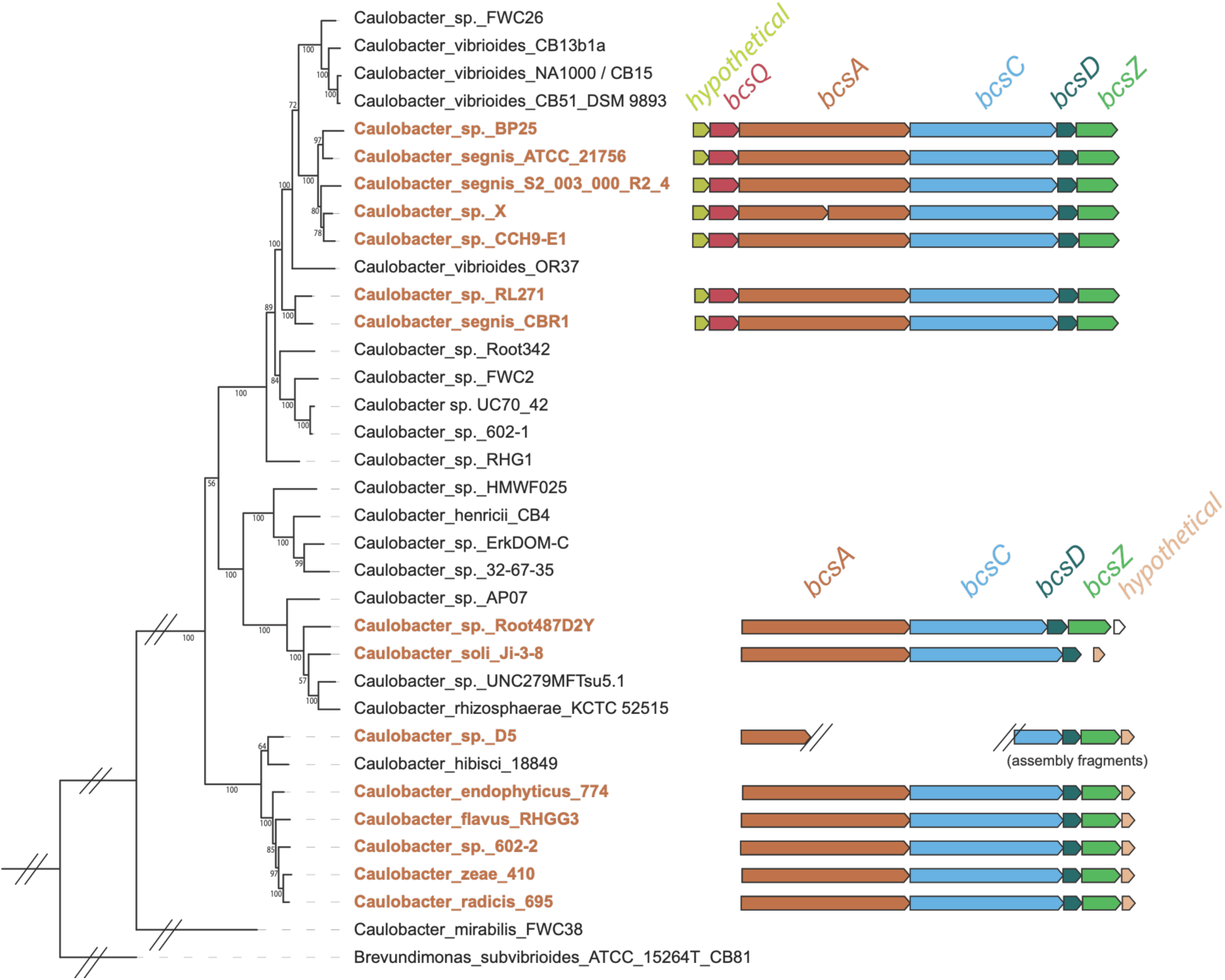
Distribution of bacterial cellulose synthesis (*bcs*) gene systems in *Caulobacter*. Two different *bcs* loci were identified in the genomes of the isolates presented in Figure 1. Genomes encoding *bcs* genes are highlighted in bold and brown font, with the corresponding *bcs* locus schematized on the right. The “hypothetical” genes, which are presumably accessory factors, are similar within each group, but distinct between groups. The phylogenetic tree is the same as in Figure 1. *bcs* genes were not identified in the genomes of the isolates in black text.

